# Insights into optimization of oleaginous fungi – genome-scale metabolic reconstruction and analysis of *Umbelopsis* sp. WA50703

**DOI:** 10.1101/2024.10.01.616082

**Authors:** Mikołaj Dziurzyński, Maksymilian E. Nowak, Maria Furman, Alicja Okrasińska, Julia Pawłowska, Marco Fondi

**Author notes:** Address correspondence to Mikołaj Dziurzyński.

## Abstract

Oleaginous fungi, known for their high lipid content—up to 80% of their dry mass—are of significant interest for biotechnological applications, particularly in biofuel and fatty acid production. Among these, the genus *Umbelopsis*, a common soil saprotroph of the Mucoromycota phylum, stands out for its rapid growth, low nutritional requirements, and ability to produce substantial amounts of lipids, especially polyunsaturated fatty acids (PUFAs). Despite previous studies on lipid production in *Umbelopsis*, metabolic engineering has been underexplored. This study fills that gap by presenting the first comprehensive metabolic model for *Umbelopsis* sp. WA50703, encompassing 2413 metabolites, 2216 reactions, and 1629 genes (*i*MD1629). The model demonstrated strong predictive accuracy, correctly predicting metabolic capabilities in 82.1% of cases when evaluated against experimental data. Using the Flux Scanning based on Enforced Objective Flux (FSEOF) algorithm, the study identified 33 genes linked to 23 metabolic reactions. Notably, reactions catalysed by acetyl-CoA carboxylase and carbonic anhydrase emerged as prime candidates for up-regulation. These findings provide a solid framework for future metabolic engineering efforts to optimize PUFA production in *Umbelopsis* strains.

**Importance:** *Umbelopsis* strains are capable of producing valuable compounds like polyunsaturated fatty acids (PUFAs). These compounds are essential for human health, found in various foods and supplements that support heart and brain function. In this study we developed a computer model to better understand how this fungus works at a metabolic level, guiding further research investigations towards optimization of PUFAs production in a cost-effective way. This research lays the groundwork for future innovations in metabolic engineering of *Umbelopsis* PUFA production leading to healthier food options and a more sustainable food system, directly impacting everyday life.

## Introduction

Oleaginous fungi are organisms notable for their high lipid content, which can constitute up to 80% of their dry mass (1). They belong to different taxonomic groups and can be both filamentous fungi and yeasts. In their natural habitats, the lipids primarily serve as the main energy storage material, but they also play critical roles in processes such as symbiosis stabilization (2, 3). The ability of oleaginous fungi to accumulate large amounts of lipids has led to extensive studies on their potential for biotechnological applications, particularly in the production of biofuels and valuable fatty acids (4, 5).

Members of *Umbelopsis* genus are common soil saprotrophs belonging to the Mucoromycota phylum, with global distribution, and predominantly associated with forest soils (6, 7). Known for its relatively rapid growth rate and low nutritional requirements, members of *Umbelopsis* have garnered significant interest in biotechnological studies focused on lipid production, particularly on polyunsaturated fatty acids (PUFAs) (8). Although lipid production of some *Umbelopsis* strains is well studied, the phylogeny of the genus is still not resolved, resulting in several misnaming and classification biases (9). The ability of *Umbelopsis* fungi to produce high lipid content, especially when cultured with glucose, may reach up to 74% of dry cell weight (10). Valuable PUFAs such as linoleic acid (Δ9,12 C18:2) and gamma-linolenic acid (Δ6,9,12 C18:3) were shown to constitute up to 18.8% and 5.1% of the lipid mass, respectively (11, 12).

While lipid production optimization efforts for members of *Umebelopsis* genus have primarily focused on growth media and general culture parameters, metabolic engineering approaches have been notably absent. Unlike other oleaginous fungi, which have been extensively analysed using bioinformatics to identify potential gene optimization targets (5, 13, 14), *Umbelopsis* sp. has not undergone such detailed investigations.

In depth genomic engineering analyses from other oleaginous fungi have highlighted long-chain fatty acid desaturases, elongases, and NADPH-generating reactions like the malic enzyme reaction as key targets for optimization (5, 13, 15). Although similar potential interventions have been suggested for *Umbelopsis* (8), they have not been thoroughly explored, partially due to the limited availability of comprehensive genomic and experimental data necessary to guide such efforts. However, reconstructing and analysing the organism’s metabolic model could address this gap, providing a valuable framework for identifying and exploring metabolic engineering targets for future.

Metabolic models have a wide range of applications, including optimizing product synthesis, discovering novel reactions and metabolic pathways, analysing the evolution of metabolic networks, and describing the robustness of metabolic systems (16–18). In the fungal kingdom, they have been extensively used to understand cellular adaptation to a wide array of environmental and genetic perturbations, optimize production of various compounds, identify novel transporter proteins, and even to study the metabolic determinants of the Crabtree effect (19–22).

This work presents the first comprehensive reconstruction of the metabolic model of *Umbelopsis* sp. WA50703. The model has been rigorously validated against available experimental data and examined for potential gene modifications to enhance the production of polyunsaturated fatty acids (PUFAs). Using the Flux Scanning based on Enforced Objective Flux (FSEOF) algorithm, multiple optimization strategies were identified, highlighting two critical reactions associated with the initiation and elongation of fatty acid synthesis as primary targets for metabolic engineering.

## Materials and Methods

### 1. Genome assembly and annotation

Raw genome sequencing data of *Umbelopsis* sp. WA50703 has been downloaded from the GenBank database (accession id: SRR26874108). The reads have been subjected to quality control and trimming using fastp v0.22.0 (23). Next, the reads were assembled using SPAdes v3.15.5 assembler with *isolate* flag (24). The assembly’s quality has been assessed using QUAST v5.2.0 run with *fungus* flag (25). Additionally, BUSCO v5.4.2 (run with mucoromycota_odb10) and CheckM2 were used to check assembly completeness and putative assembly contamination (26, 27). Taxonomy has been confirmed by aligning ITS sequences extracted from the assembly using ITSx v. 1.1.3 versus UNITE database (28, 29).

Structural and functional annotation was conducted using the funannotate pipeline v. v1.8.14 (30). Briefly, repeat-masking was performed using TANTAN software (31). Masked contigs were subjected to gene calling, which was conducted using multiple programs: Augustus, GeneMark, GlimmerHMM and SNAP supplied with RNAseq evidence (European Nucleotide Archive run ID: SRR13866027) to improve prediction accuracy (32–35). Results obtained from all programs were integrated using Evidence Modeler v. 1.1.1 (36). tRNAs were predicted using tRNAscan-SE v. 2.0.9 (37). Predicted coding sequences were annotated using “annotate” script included in the funannotate pipeline. The script annotates coding sequences using multiple reference databases, such as Pfam, UniProt CAZy, MEROPS, and BUSCO groups (38–41). Additionally, the pipeline was supplied with InterProScan and EGGnogMapper results to integrate into the annotation (42, 43). Finally, the pipeline performed protein cellular localization annotation based on SignalP results and annotation of biosynthetic gene clusters based on antiSMASH results (44, 45). Finally, the annotation was manually curated using MAISEN web application (46).

### 2. Draft model reconstruction and manual curation

The draft model was *de novo* reconstructed using Pathway Tools v. 26.5 (47). This software employs a multi-stage process to identify metabolic reactions through genome annotation word-matching and protein sequence alignment with the MetaCyc database. It also performs a basic gap-filling process to complete metabolic pathways by adding potentially present but previously unidentified reactions. The initial draft reconstruction was then manually curated following the procedure outlined by Thiele and Palsson (48). This curation process included evaluating the feasibility of biomass precursor production, aligning the model with SBML standards, verifying reaction directionality, incorporating the mitochondrion compartment, and correcting erroneous CO₂ assimilation and energy generation reactions.

The biomass reaction was defined using a spreadsheet developed by Ye et al. in their work on the *M. alpina* model (13). Macromolecular formulations were supplemented with data from the available literature. DNA, RNA, and amino acid contributions were calculated using the BOFdat algorithm, while lipid and carbohydrate fractions were adjusted based on experimental data from multiple studies (49).

### 3. Experimental data collection

The available literature was thoroughly reviewed for experimental data pertaining to growth of *Umbelopsis* sp. in various conditions. Special attention was paid to culture experiments measuring fungal growth coupled with depletion of carbon substrates, and experiments testing utilization of different carbon sources by *Umbelopsis* members.

In the first case, five articles, with a total of seven different experimental datasets, have been identified. Available growth curves and carbon source depletion curves were extracted and subjected to further calculations. Growth curves were standardized using growthcurver v. 0.3.1 package run in RStudio v. 4.2.1 environment (50). Maximal growth rates and carbon source uptake rates were calculated as described by Fondi et al (51).

Carbon assimilation data has been extracted from the work of Pawłowska et al (52). The data included Biolog phenotypic array results run with *U. isabellina* CBS167.80. The data covered tests run with 95 different carbon sources, each with three replicates. Due to high data variability, positive assimilation capability was assumed only when growth was observed in at least two out of three replicates.

### 4. *In silico* optimization of lipids production

The Flux Scanning based on Enforced Objective Flux (FSEOF) algorithm was utilized to identify potential bottlenecks and target genes for optimizing the biosynthesis of long-chain fatty acids (53). A total of 59 model versions, each with a different carbon source, were subjected to FSEOF analysis. The analysis was performed using the RAVEN Toolbox 2, version 2.8.4, run within the MATLAB R2023 environment (54, 55). The configurations specified ’Biomass_reaction_1’ as the biomassRxn parameter and ’DM_linoleate_c’ as the targetRxn parameter. Results from all tested models were aggregated and analyzed, focusing on reactions that appeared in at least five solutions and had an FSEOF slope value exceeding 2, thereby identifying the most promising reactions.

### 5. Data availability

The *i*MD1629 model, along with custom tests is available under the following URL: https://github.com/mdziurzynski/iMD1629_uisabellina_MM

## Results

### 1. Genome assembly and annotation results

Quality assessment of the raw sequencing data indicated high data quality, with over 99% of reads passing all filtering steps. The assembled genome of *Umbelopsis* sp. WA50703 comprised 234 contigs with a mean sequencing depth of 134x and an average GC content of 41.95%. The L50 and N50 metrics were 10 and 750,386, respectively (Supplementary Table 1). BUSCO analysis revealed 95% genome completeness, with 2.3% of genes fragmented and 2.7% missing. CheckM2 showed genome completion at 89.43% with 8.63% possible contamination. Species level identification enable to classify the strain within *U. isabellina* section. However, the strain does not form monophyletic group with *U. isabellina* type material, therefore, we refrain from using species name for *Umbelopsis* sp. WA50703 strain.

During genome annotation, 8,693 genes were identified, including 8,548 encoding mRNAs and 145 encoding tRNAs. A total of 7,271 genes were annotated with at least one functional domain, and detailed protein functions were assigned to 2,832 of them. Additionally, 1,464 proteins were annotated with an EC number. Overall, only 1,277 proteins remained without any functional annotation. Detailed annotation results are available in Supplementary Table 1.

### 2. Model description

The reconstructed model was named iMD1629 and consisted of 2203 reactions, 2404 metabolites and covered 1629 genes. The model is highly annotated, as over 75% of all reactions, and nearly 100% of metabolites were annotated with an appropriate BioCyc identifier. Beside BioCyc, metabolites and reactions were also annotated with identifiers to the following classifications and databases: Enzyme Commission number, Chemical Entities of Biological Interest (chEBI), International Chemical Identifier (InChl), PubChem, KEGG, Rhea, Chemical Abstracts Service number (CAS) and others.

The model consists of three major compartments: extracellular space, cytosol and mitochondrion. Using DeepLoc 2.0, a deep learning algorithm for predicting subcellular localizations of eukaryotic proteins, it was possible to separate reactions, and by extension the metabolites, between the compartments. There were 60, 1581 and 389 reactions in extracellular compartment, cytosol and mitochondrion, respectively. Additionally, there were 173 transport reactions. Transport reactions were examined manually, and only relevant and feasible reactions, according to the MetaCyc or the Transporter Classification Database, were added.

The biomass equation was curated using experimental data available in the literature from studies on *Umbelopsis* strains. Although exact data for DNA, RNA, and protein content were unavailable, their biomass contributions were estimated based on genome content, using a methodology similar to that employed by the BOFdat Python package (49). Data published by Muszewska et al. (2021) provided a detailed estimation of the carbohydrate fraction, predominantly composed of glucose, galactose, mannose, fucose, and N-acetylglucosamine (7).

Since most experimental data regarding *Umbelopsis* focus on the organism’s lipid production under various medium and environmental conditions, the lipid fraction was curated with the highest accuracy and detail (8, 9, 56). The data enabled a comprehensive partitioning of the lipid fraction into neutral lipids and phospholipids, along with detailed contributions from seven different fatty acids and five classes of phospholipids. However, because the lipid data derived from highly optimized cultures, the exact contributions of each compound were adjusted for a minimal medium environment. A full description of biomass fractions and their contributions is available in Supplementary Table 2.

The reconstructed model was evaluated using Memote, a comprehensive test suite for genome-scale metabolic models, and received an overall score of 68%. The highest scores were achieved in data annotation categories, including "Metabolite Annotation" and "Reaction Annotation," while the lowest score was in "Consistency." The complete Memote report is available as Supplementary File 1. Additionally, Memote was used to set up a GitHub repository to host the latest version of the model. Beyond the basic auto-generated Memote content, such as Memote reports, the repository also includes a set of experimental validation tests for the model. The repository can be accessed at the following link: https://github.com/mdziurzynski/iMD1629_uisabellina_MM.

### 3. Model validation on experimental data

The reconstructed model was evaluated against available experimental carbon assimilation data, extracted from the work of Pawłowska et al (52). The data contained information about the *U. isabellina* CBS167.80 growth on 95 carbon sources, i.e. 7 amines/amides, 13 amino acids, 15 carboxylic acids, 46 carbohydrates, 5 polymers, and 9 other compounds. All simulations were run with four exchange reactions, reflecting essential compounds present in *U. isabellina* minimal medium, left unconstrained. They were phosphate, sulphite, sulphate and ammonium exchange reactions. Additionally, exchanges for carbon dioxide, oxygen, water and hydrogen ions were also set open.

The analysis demonstrated an overall agreement of 82.1% between the experimental results and in silico simulations. The model accurately predicted growth on 42 carbon sources that also showed growth in vitro and correctly predicted no growth on 36 carbon sources, aligning with the in vitro results. However, there were discrepancies for 17 carbon sources: 10 showed no growth in silico, while 7 showed growth in silico, contrary to the experimental data.

The former group included D-glucuronic acid, Tween 80, α-cyclodextrin, β-cyclodextrin, dextrin, L-rhamnose, and sebacic acid. For these compounds, no transporters were detected, suggesting either transportation via unknown mechanisms or proteins, or initial extracellular degradation by unidentified secreted enzymes. Specifically, no substrate-specific enzyme responsible for the degradation of D-glucuronic acid was identified. Tween 80 likely undergoes non-specific degradation by esterases, lipases, and cutinases. The absence of cyclodextrinase (EC 3.2.1.54) in the genome assembly precluded the inclusion of cyclodextrin (both α- and β-cyclodextrin) degradation reactions in the model. Additionally, no relevant enzymes for the degradation pathways of L-rhamnose were identified, and enzymes required for the degradation of alpha, omega-dicarboxylate acids (sebacic acid) such as EC 6.2.1.23 and EC 6.2.1.10 were also absent.

The latter group included xylitol, fumaric acid, β-hydroxy-butyric acid, α-keto-glutaric acid, L-lactic acid, succinic acid, L-aspartic acid, L-proline, L-serine, and L-threonine. Most of those compounds participate in core metabolic pathways and therefore their positive and, at the same time, incorrect assimilation result most probably arises from misidentification of their respective transporter proteins. Full table comparing *in silico* simulations with experimental data results is available as Supplementary Table 3.

Biolog experimental results have also been used to evaluate overall utilization of reactions included in the model. Information about active and inactive reactions from feasible *in silico* solutions for 59 different carbon sources were pooled and analysed (Fig. 2A). The results showed that, on average, each solution contained 336 active reactions. Pooled data analysis revealed that in all the 59 different solutions 209 reactions were always active (9.5%), 445 reactions were active at least once (20.2%) and 1549 reactions were never active (70.3%) (Fig. 2C). While the “active at least once” reactions were heavily associated with supplied carbon source, with a clear division between carbohydrates and amino acids, further analysis showed interesting recurrent patterns. Out of the 444 auxiliary reactions (the “active at least once” set), 39 (8.78%) were extremely close in their activity pattern to the core reactions, as at most they were inactive in only 10% of the investigated solutions (Fig. 2D). The following 186 reactions (41.89%) could be interpreted as necessary for assimilation of different carbon source types, such as amino acids or carbohydrates. Their activity was observed in not less than 10% and no more than 90% of solutions. Finally, the remaining 219 reactions, which constituted 49.32% of the auxiliary set, were interpreted as rarely active but vital for the metabolism of specific carbon sources, as they were active in no more than 10% of solutions. Detailed pathway analysis of this latter group identified three pathways that were active exclusively when growth was simulated on very specific carbon sources. These pathways were: adenosine degradation through urate and allantoine (10 rare reactions, BioCyc IDs: SALVADEHYPOX-PWY, PWY-5691, PWY-5694), D-galacturonate degradation (5 rare reactions, BioCyc ID: PWY-6491), and L-arabinose degradation (3 reactions, BioCyc ID: PWY-5515).

**Fig. 1.**
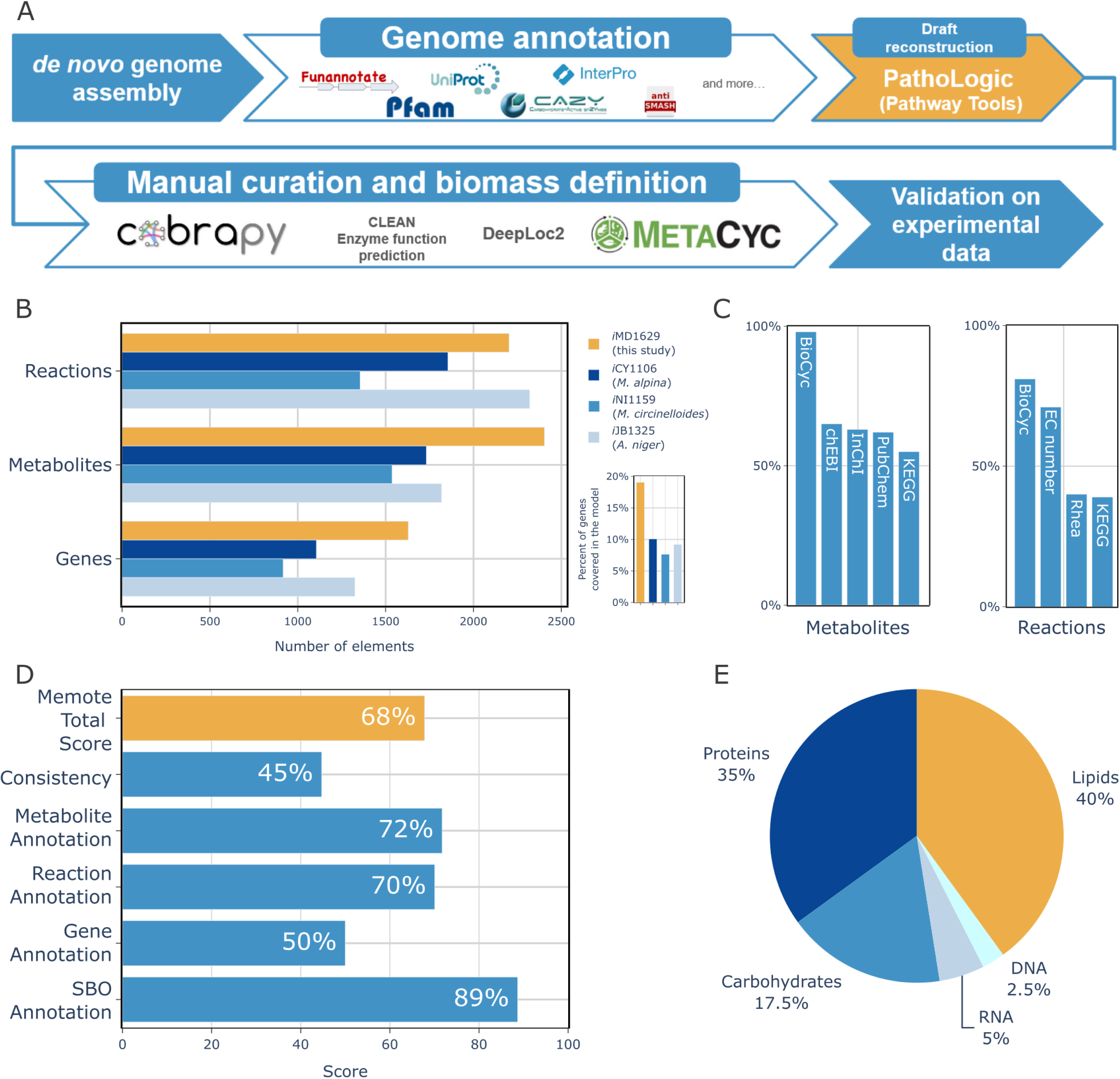
Model reconstruction pipeline and model metrics. A: main steps of the model reconstruction pipeline, B: comparison of iMD1629 reconstructed in this work with three other high-quality fungal models; C: iMD1629 reactions and metabolites annotation coverage; D: Memote results for iMD1629; E: main biomass precursors and their contributions.

**Fig. 2.**
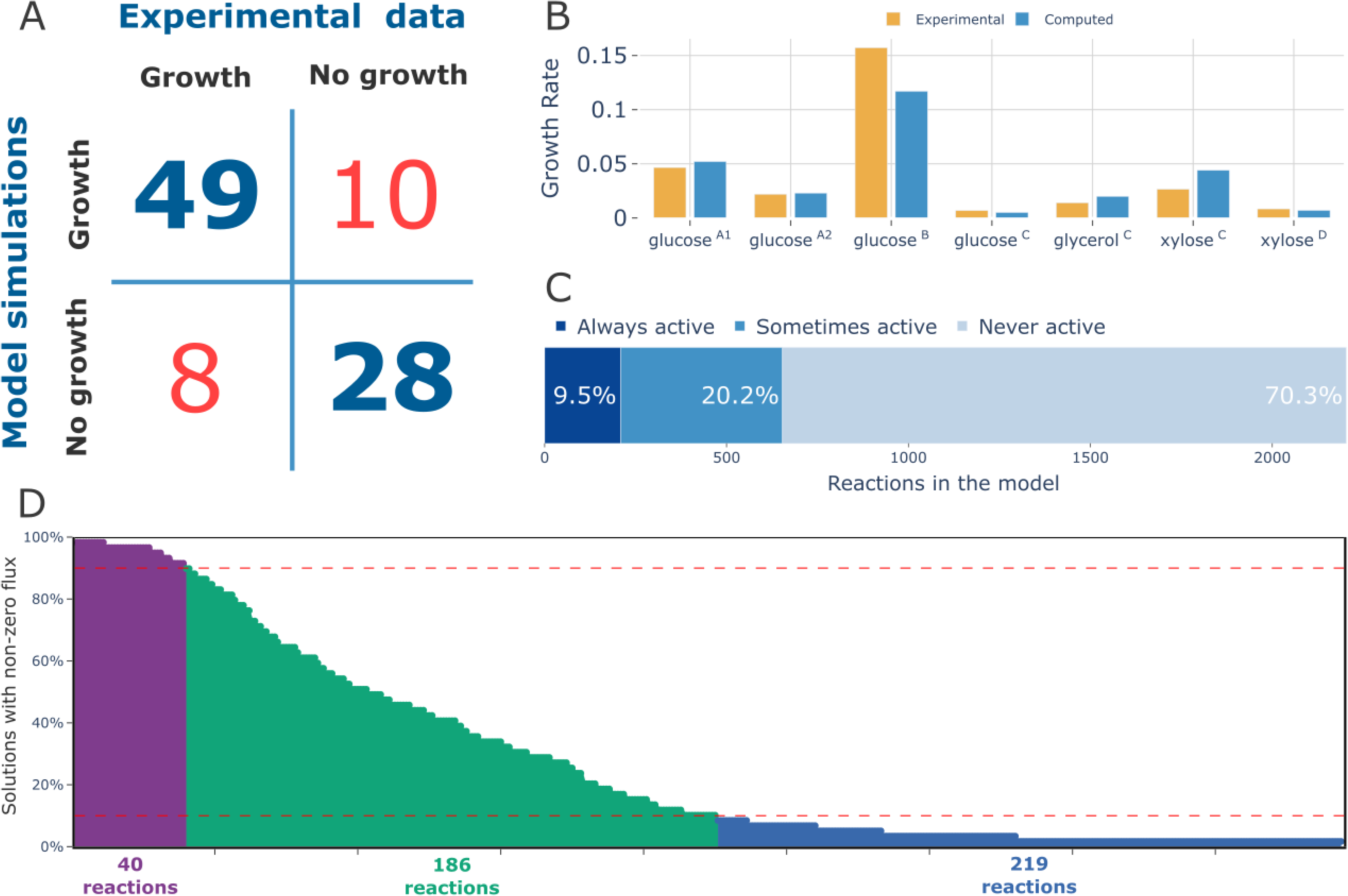
Model validation results. A: comparison of Biolog phenotypic microarray results for *U. isabellina* with *i*MD1629 model simulations; B: comparison of experimental and computed growth rates of *U. isabellina* with different carbon sources, ^A1^-flask cultures data from Chatzifragkou A. et al (10), ^A2^-cultures run in 3 litre bioreactor from Chatzifragkou A. et al (10), ^B^-cultures run in 3 litre bioreactor from Meeuwse P. et al (57), ^C^-flask cultures data from Fakas S. et al (58), ^D^-flask cultures data from Gardeli C. et al (11); C: utilization of model reactions observed in simulations run with different carbon sources, D: activity of “Active at least once” reactions set among all 59 simulations with different carbon sources. Red dashed lines separate 10% and 90% activity thresholds, dividing the graph in three groups of near core reactions (violet), carbon source type related reactions (green), carbon-source specific reactions (blue).

Further, we also evaluated the capability of the reconstruction to quantitatively resemble the growth features of *Umbelopsis sp*. Accordingly, growth rates predicted by the model were compared against growth rates extracted from growth curves available in the literature. Although the experimental data came from experiments run with different culture conditions and even different *U. isabellina* strains, the results showed high similarity between *in silico* and *in vitro* growth rates. Experimental data covered four experiments run with glucose as the main carbon source, two with xylose and one with glycerol. Fig. 2B presents growth rates from each experiment and their computed pair from the model simulations. Overall, the model showed remarkable accuracy in predicting the growth rates of *U. isabellina* on known carbon sources. Only in two out of seven comparisons, the difference was higher than 30%, as experimental growth rate was 1.3 higher and 0.6 lower in case glucose-B and xylose-D experiments. In all the other cases, the simulation results and experimental values resembled each other (Fig. 2B). Overall, these data suggest that the iMD1629 reconstruction can reliably predict *U. isabellina* growth phenotypes both qualitatively and quantitatively.

### 4. Lipid production optimization – case study

The reconstructed model has been employed for identification of gene amplification targets for improvement of long fatty acids production. The analysis was conducted using FSEOF algorithm (53). FSEOF performs a series of model simulations, slowly shifting model objective from biomass production to a desired by-product. This approach allows identifying reactions that correlate with increased shift towards production of intended by-product.

FSEOF analysis showed 297 potentially interesting gene targets, most of which were reported for only one specific carbon source. To identify the most important and the most prevalent gene targets, final reaction set was filtered to contain reactions that were reported for at least 5 different carbon sources. Additionally, a FSEOF slope threshold of greater than 2 was applied to ensure that only the reactions with the highest impact were retained. Fig. 3 presents a heatmap describing this set of reactions and includes 23 reaction instances retrieved after simulations with 56 different carbon sources. There were 5 reactions that showed up in nearly all solutions as potential targets for improvement. Two of them, ACETYL-COA-CARBOXYLTRANSFER-RXN and RXN0-5224, were associated with fatty acids biosynthesis initiation and elongation. The other three, CoA transport (c -> m), PYRUVDEH-RXN and Pyruvate transport (c -> m), were associated with transport and preparation of main TCA cycle substrates. The next six reactions were associated with TCA cycle and introduction of L-glutamate into the TCA cycle. These reactions were reported mainly in solutions obtained for growth simulations with amino acids and amides/amines set as main carbon sources. The last and, at the same time, the biggest set was composed of 12 reactions, 11 of which constitute the core of glycolysis. As expected, those reactions were reported mostly in growth solutions run with carbohydrates as main carbon sources.

**Fig. 3.**
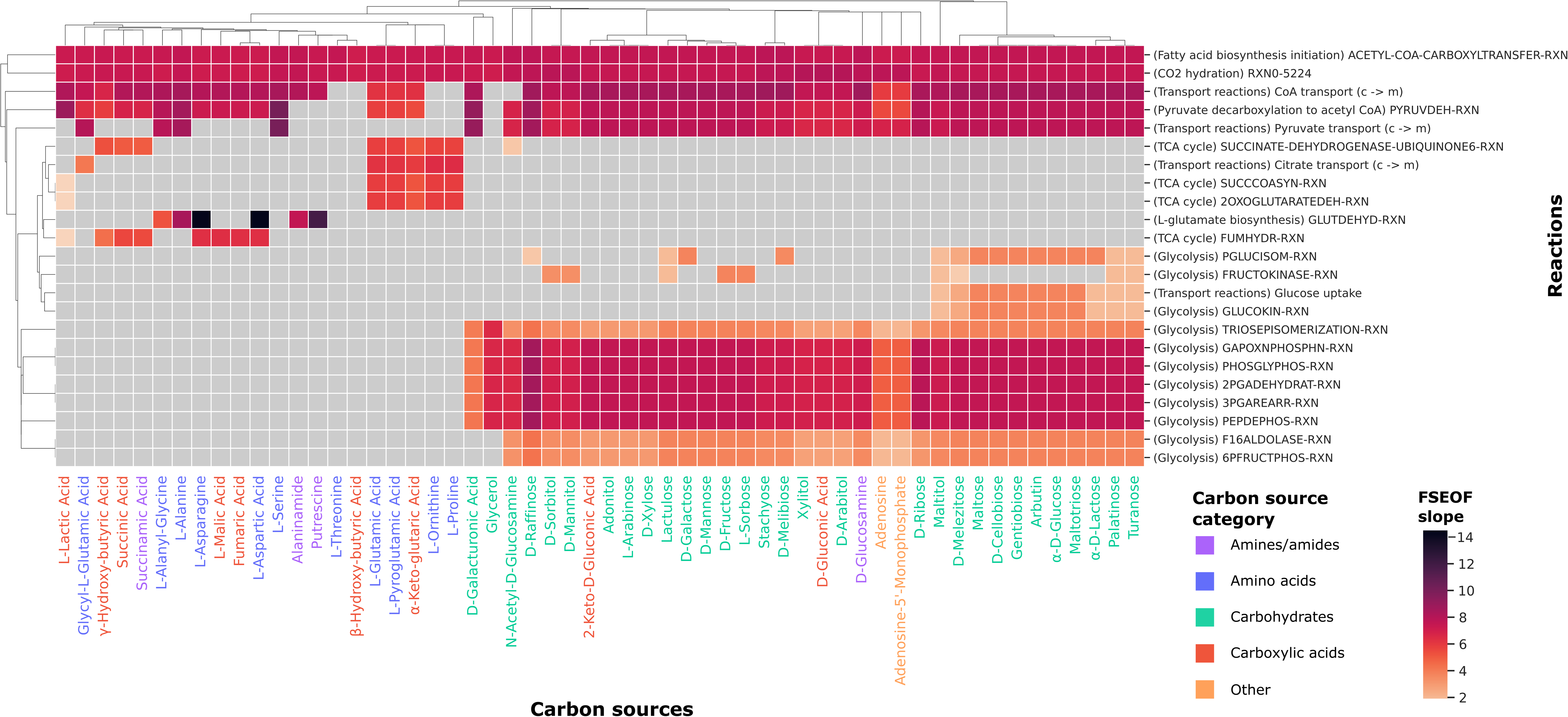
Heatmap depicting FSEOF analysis results. Slope values indicate level of co-upregulation of given reaction with an increase in production of linoleate. Grey tiles indicate that a given reaction was not identified as potential target for improvement for a given carbon source. Other than transport reactions, reaction names are MetaCyc reaction identifiers.

The set of 23 reactions highlighted by FSEOF are encoded by 33 different genes, some of which encode multi-protein complexes (Supplementary Table 4). Three of the reactions lacked gene annotation and they were “Pyruvate transport (c -> m)”, “Citrate transport (c - > m)” and “FRUCTOKINASE-RXN”.

## Discussion

High biotechnological potential of different *Umbelopsis* strains has been investigated for over 20 years, however the primary focus of these analysis was optimization of exogenous factors, such as culture conditions, for the production of various value-added compounds, predominantly PUFAs(8). Although the organism exhibits exceptional oleaginicity and grows well on different, low-cost carbon sources, its further and more detailed optimization projects are substantially hampered by the limited availability of suitable genetic toolkit and *in silico* insights that could guide such efforts.

In this study we present the first *in silico* metabolic model of *Umbelopsis* sp. WA50703 In this reconstruction high attention was paid to both genome and model annotation, in order to increase the model interoperability and utility. Genome assembly analysis agreed with the expected genome size of 22.74 Gbp, comparable with other *Umbelopsis* genome assemblies available in the GenBank database. Although BUSCO analysis yielded a high, 95% genome completeness result, CheckM2 contamination analysis showed 8.63% contamination. Further analysis using BlobTools2 showed that the contamination was detected probably due to the incompleteness and incorrectly annotated fungal sequences in the reference databases, as fungal genomes sequencing lags significantly behind other organism groups (59).

Detailed genome annotation resulted in exceptionally high genome coverage in the *i*MD1629 model (20% of all genes were present in the model), when compared with other, high-quality fungal models (between 7-10% of genes) (13, 60). Although the model contains a similar number of metabolites and reactions compared to other models, it has been reconstructed with only three compartments: extracellular, cytosolic, and mitochondrial. Eukaryotic models typically include additional compartments, such as the periplasmic space, peroxisome, nucleus, or Golgi apparatus. However, in the case of *i*MD1629, this further division of reaction space was omitted due to the lack of experimental data needed to accurately assign reactions to these compartments. Without data on inter-compartmental transport capabilities, incorporating additional compartments would result in artificial and unreliable pathway structures and flux flows that may not accurately reflect biological reality. It is important to note that, should there be a reason and sufficient data to add new compartments in the future, the model’s reaction and metabolite annotations are designed to facilitate this task.

While the experimental validation results showed a high degree of accordance with model simulations, they also highlighted significant areas for improvement. The experimental data collected from publicly available resources for *Umbelopsis WA50703* proved invaluable for model curation and testing. However, to eliminate inconsistencies between experimental results and simulations, future experiments must be conducted in a more strain-specific manner as it was done with *Aspergillus niger* by Brandl J. et al (60). Variability in growth rates obtained from glucose growth experiments can be attributed to substantial differences in experimental setups.

For example, the experiment with the highest growth rate had several differing conditions: a much higher initial concentration of nitrogen source (20 mM vs. 4 mM), a pre-culturing step to adapt the fungus to culture conditions, active aeration, larger volumes (2000 ml vs. 50 ml), and used a different *U. isabellina* strain (CBS 194.28 vs. ATHUM 2935) (57, 58). Additionally, it is important to note that *Umbelopsis* has only recently been recognized for its facultative yet persistent endohypal bacteria, which can significantly expand its metabolic capabilities bacteria (61). Therefore, before generating experimental data, it should be confirmed that the strain in use is free of these symbionts.

The model validation has confirmed its readiness for guiding metabolic engineering strategies. In this study we employed FSEOF algorithm to identify reactions that should be up-regulated in order to increase production of PUFA. The results showed clear distinction between optimization approaches for different carbon source types. Reactions included in glycolysis were characteristic for carbohydrate carbon sources, while TCA cycle reactions were characteristic for amino acid-based media. However, the analysis also yielded five reactions, that were common to nearly all carbon substrates. Two of them were simple, cytosol-to-mitochondrium transport reactions of coenzyme A and pyruvate. The second pair, ACETYL-COA-CARBOXYLTRANSFER-RXN and RXN0-5224, was associated with cytosolic fatty acids biosynthesis initiation. ACETYL-COA-CARBOXYLTRANSFER-RXN is catalysed by acetyl-CoA carboxylase (EC number: 6.4.1.2) and performs an ATP-dependent carboxylation of acetyl-CoA to malonyl-CoA, a crucial compound for fatty acid biosynthesis initiation and elongation (62). RXN0-5224, is catalysed by carbonic anhydrase (EC number: 4.2.1.1) which is a metalloenzyme responsible for reversible interconversion between CO2 and bicarbonate (63). Bicarbonate produced in this reaction is used by acetyl-CoA carboxylase for carboxylation of acetyl-CoA. Both of those reactions have been previously and successfully investigated for increasing lipid production in bacteria, algae and fungi. Although employed methods varied, from introduction of a strong, constitutive promoters for native genes, to introduction of multiple heterologous genes, their efficiency relied heavily on the target organism (64, 65). *E. coli* showed very high 6-fold increase in fatty acid production, *Yarrowia lipolytica* (2.93-fold increase) and green algae *Chlamydomonas reinhardtii* (1.16-fold) (66–68).

The last reaction was mitochondrial PYRUVDEH-RXN which is catalysed by pyruvate dehydrogenase complex (EC number: 1.2.1.104). This reaction combines pyruvate with coenzyme A into acetyl-CoA with the release of NADH. While it plays a major role in feeding carbon to the TCA cycle in the mitochondrion, metabolic engineering efforts focused on its implementation as a part of cytosolic, multistep, synthetic pathway that converts pyruvate to acetyl-CoA while producing NADPH to boost lipid biosynthesis (69).

Therefore, all three of those reactions, and essentially their associated genes, may be a viable, first choice targets for up-regulation in *Umbelopsis* WA50703 for the production of PUFAs.

## Conclusions

In this work the metabolic model of oleaginous fungi *Umbelopsis* sp. WA50703 has been reconstructed. The model was reconstructed with special attention paid to metabolites, reactions and genes annotation quality. Experimental validation of the model showed 82.1% concordance with phenotypic microarray data and 68% Memote Total Score. A case study analysis using the model, aiming at identification of possible targets for improvement to increase PUFAs synthesis yielded 23 reactions, out of which three have been identified as the most promising ones as they were identified on nearly all tested carbon sources. The model provides a promising starting point for guided metabolic engineering of *Umbelopsis* and deeper investigations into its metabolic potential.

